# Comparison of different methods for average anatomical templates creation: do we really gain anything from a diffeomorphic framework?

**DOI:** 10.1101/277087

**Authors:** Vladimir S. Fonov, D. Louis Collins

## Abstract

In the field of computation anatomy, the diffeomorphic framework is widely used to perform analysis of human brain anatomy in both healthy and diseased populations. While useful for analysis, the framework imposes certain implementation constraints that do not necessarily result in improved accuracy of inter-subject co-registration in case of average anatomical template (AAT) construction – a common technique used in large population studies. In this work, we evaluated several state-of-the-art non-diffeomorphic and diffeomorphic non-linear registration frameworks in terms of their ability to build AATs. While all methods generated well behaved transforms, we found that the diffeomorphic framework does not automatically guarantee an increase of accuracy in average anatomical template construction.

## 1 Introduction

Average anatomical templates (AAT) are useful tools in computational anatomy (Mazziotta, Toga et al. 1995). They are applied in a wide variety of methods requiring registration to a common space for comparison, for example when performing Voxel-Based Morphometry (VBM) analysis, serving as priors for the automatic tissue classification and automatic anatomical structure segmentation. Multiple strategies for creating AATs have been created during the last 19 years. Starting with voxel-wise arithmetic averaging of manually registered scans (Evans, Collins et al. 1993), automatic linear co-registration (Collins, Neelin et al. 1994) and then non-linear average templates using elastic registration (Guimond, Meunier et al. 1998), free-form deformation with B-Splines (Bhatia, Aljabar et al. 2007), and large deformations diffeomorphic mapping (LDDM) (Lorenzen, Davis et al. 2005). Several studies testing the impact of template construction have been published (Wang, Seghers et al. 2005), (Klein, Ghosh et al. 2010), (Avants, Tustison et al. 2011) with overall conclusion that templates based on the average shape and intensity perform better in anatomical structure segmentation then direct pair-wise registration.

Among the various currently available methods, one important distinction appears: some methods are based on a diffeomorphic framework (Avants, Tustison et al. 2011), (Lorenzen, Davis et al. 2005), while others aren’t (Guimond, Meunier et al. 1998), (Bhatia, Aljabar et al. 2007). The diffeomorphic template building framework guarantees that the generated transforms are smooth, invertible, bijective maps between all volumes, such that the forward and inverse transforms are differentiable. This preserves object topology through the transform, and allows using methods developed in the field of differential geometry and generally facilitates analysis. While in theory the use of a linear elastic regularizer (e.g., (Collins, Holmes et al. 1995)) or arithmetic operations on vector deformation fields do not guarantee a diffeomorphic transformation, in practice this is often the case because of the constraints implemented in optimization and regularization; the resulting transformations are smooth, invertible and both forward and inverse transforms are differentiable numerically. Some of these constraints are not unlike those needed for numerical implementation of a diffeomorphic mapping on a discrete grid. While some researchers view the diffeomorphic framework as a kind of silver bullet, able to automatically improve and/or justify results of statistical morphological studies of human brain anatomy, how do these imposed constraints affect average anatomical template construction? Are diffeomorphic registration methods really needed?

The purpose of this study is to evaluate if the diffeomorphic framework guarantees better results in the limited scope of building average anatomical templates, where *better* is defined in the sense of improving co-registration accuracy between subjects chosen to create the template. In this paper, five algorithms based on the publicly available tools are compared, with the ANTS method (Avants, Tustison et al. 2011) and simple linear average serving as a baseline reference. In the spirit of reproducible research, the source code for these comparisons is made publicly available for at the url: http://anonymous.web.site [<u>note: anonymous only during review</u>].

## 2 Materials and Methods

In this study, two databases are used. First, T1-weighted (T1w) MRI data from 152 normal subjects from the International Consortium for Brain Mapping (ICBM) project (1mm thick slices, TR=18 ms, TE=10 ms, flip angle 30° with 1mm in-plane resolution, acquired on a 1.5T Philips Gyroscan scanner (Mazziotta, Toga et al. 1995)); and second, MRI scans and manually segmented labels of 40 normal subjects from Probabilistic Brain Atlas (LPBA40) at the Laboratory of Neuro Imaging (LONI) at UCLA (Shattuck, Mirza et al. 2008) (High-resolution 3D Spoiled Gradient Echo (SPGR),1.5-mm slices, TR range 10.0–12.5 ms; TE range 4.22–4.5 ms; in-plane voxel resolution of 0.86 mm or 0.78 mm, acquired on a GE 1.5T scanner).

Anatomical scans from ICBM project were corrected for image intensity inhomogenieties (Sled, Zijdenbos et al. 1998), linearly registered to MNI152 stereotaxic space (Collins, Neelin et al. 1994), the intensity range was linearly normalized to [0, 100] using histogram matching and brain extraction was performed using BET (Smith 2002). Scans in “delineation space” from the LPBA40 database were already processed for intensity inhomogeneity and stereotaxic registration, and were only normalized to be in the [0,100] interval and used together with the manually-defined anatomical labels describing 55 structures.

Scans from both databases were automatically segmented into three brain tissue classes: white matter (WM), gray matter (GM) and cerebro-spinal fluid (CSF) using unsupervised finite mixture model classifier (Tohka, Krestyannikov et al. 2007) which does not require usage of spatial priors. These tissue labels (and the anatomical labels of the LPBA40) are used below in the goodness of fit metrics used to compare the different registration strategies.

### Minimum deformation template

The algorithm to build a minimum deformation template is similar to that described in (Avants, Tustison et al. 2011) (in the script buildtemplateparallel.sh available in ANTs source code) and in (Fonov, Evans et al. 2011). The strategy used here is formulated as the following:

1. For a given template *Ti-1* and subjects *Ik* (where *k=1‥K*), find (non-linear) transformations *Si,k* minimizing ***D****(Ti-1,Ik(Si,k))* for each subject *k*, where ***D*** represents the costfunction used, optionally using *Fi-1,k*, the non-linear transformation from the previous iteration for initialization.
2. Estimate the Frechet mean, *Zi*,, in the space of transformations, minimizing 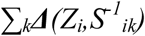 where ***Δ*** represents a distance function in the space of non-linear transformations, and 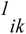 is the inverse of *Sik*, such as 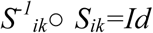.
3. Calculate *Fik=****U****(Sik,Zi)*, where ***U*** is the update rule.
4. Estimate new template *Ti*, minimizing *Σ*_*k*_***D****(Ti,Ik(Fik)).* In the case when D is cross-correlation or sum of squared differences, the arithmetic mean is the solution: *Ti=1/K·ΣkIk(Fik)*,
5. Repeat steps 1-4 until convergence

It is possible to adopt above algorithm to use different non-liner registration schemes, both Diffeomorphic and Elastic, and in the current study we have evaluated performance of the following template building methods:

- ANIMAL (Collins, Holmes et al. 1995) elastic non-linear registration algorithm with correlation-coefficient intensity cost function ***D***, and sum of squared differences distance ***Δ*,** resulting in the Zi=1/K*ΣkSik*,and update rule *U(x,y)=x○y.*
- Diffeomorphic Demons (DD) (Vercauteren, Pennec et al. 2009) with symmetrized gradients used to compute the demons forces, diffeomorphic non-linear registration algorithm, sum of squared differences cost function ***D***, sum of squared differences distance ***Δ*,** and update rule *U(x,y)=x○y.* Implemented in ITK 3.20.1 (Ibanez 2005).
- Thirion’s Demons or Non-Diffeomorphic Demons (NDD), with symmetrized gradients (Vercauteren, Pennec et al. 2009), sum of squared differences cost function ***D***, sum of squared differences distance ***Δ*,** and update rule *U(x,y)=x○y.* Implemented in ITK 3.20.1 (Ibanez 2005).
- Log-space Diffeomorphic Demons (LDD) (Vercauteren, Pennec et al. 2008) with symmetrized gradients, in this case all operations on the transformations are performed in log-domain on velocity fields *Vik*, which can be converted into a diffeomorphic transformation via exponential mapping, i.e *Sik=exp(Vik)*. In this case, we use sum of squared differences cost function ***D***, sum of squared differences distance ***Δ*** calculated in log-domain and update rule U(x,y)=x+y also calculated in log-domain.
- Symmetric group-wise normalization template building algorithm (Avants, Tustison et al. 2011) as defined in script buildtemplateparallel.sh in the ANTs package (ANTs) with the exception of disabling intensity normalization and sharpening (Laplaci-anSharpeningImageFilter) filter inside AverageImages tool, using cross-correlation intensity cost-function.
- Straight linear averaging (LINEAR), i.e., without any non-linear registration.

### Hierarchical processing

Almost all the template building algorithms described above (apart from LINEAR) use a hierarchical approach for varying parameters of non-linear registration, starting with a rough estimation of deformation (or velocity) fields and progressively increasing resolution. We have adopted a scheme compatible with the one published in (Fonov, Evans et al. 2011): for the ANIMAL method: 4 iterations at 32mm resolution of deformation field, 4 at 16mm, 4 at 8mm, 4 at 4mm and 4 at 2mm with 20 iterations in total; for DD,NDD and LDD we have started at 4 iterations at 16mm resolution, 4 at 8mm, 4 at 4mm, 4 at 2mm and 4 at 1mm; for ANTs we have used the scheme published in (Avants, Tustison et al. 2011) – 3 iterations with 30×50×0 registrations steps and 3 iterations with 50×90×20 registration steps. All template building algorithms evaluated here produce a dense deformation field for each subject, enabling it to be warped into the space of the final AAT. These deformation fields may be applied to the tissue classification maps and anatomical labels maps to produce probabilistic atlases of corresponding tissue types and anatomical structures.

### Performance metrics

The relative performance of each algorithm is evaluated using the following metrics:

- Generalized Tannimoto Coefficient (GTC, (Crum, Camara et al. 2006)). GTC is applied to the tissue and anatomical label maps resampled using nearest-neighbour interpolation using the final non-linear deformation fields obtained during template construction. Described by Eq.1, which is generalization of Tannimoto or Jaccard overlap coefficient in the case of fuzzy labels A_kli_,B_kli_, multiple labels and multiple subjects. 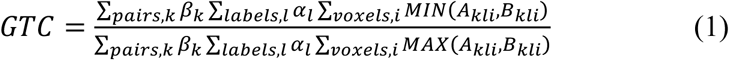 By collapsing summations depending on the situation, it is possible to apply this formula in different cases (e.g., voxel-wise across all subjects, region-wise between only two subjects, etc…) To evaluate the accuracy of each AAT building algorithm, the GTC was applied on both a voxel-wise level for the tissue-classification and anatomical structures maps warped into the common space, and overall for the whole brain. Also, we have calculated both overall GTC for all subjects participating in AAT building and for all possible pairs to estimate the range of overlap values.
- Harmonic Energy (HE, (Vercauteren, Pennec et al. 2009)). HE is applied to the individual deformation fields, described in Eq. 2, where J is Jacobian matrix, V is the total number of voxels and ||·|| is Frobenius norm. The HE represents the degree of smoothness of the transform. For equivalent GTC, smaller HE is better. 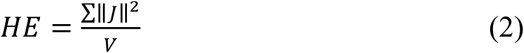
- Sharpness (Sh). Sh is applied to the result of template building process, determined by Eq. 3 where G is Gaussian smoothing kernel with a full-width half-max (FWHM) of 1mm. *Sh* represents relative visual sharpness of the image *I*.

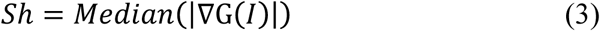

### Parameter optimization

Demons-based algorithms have two tunable parameters: deformation field smoothing sigma *s* and update field smoothing sigma *g*, to control the degree of regularization in the non-linear registration process. The ICBM152 dataset was used to find the optimal values of *s* and *g* for the LDD template building algorithm.

## 3 Results

### Parameter optimization using ICBM 152 dataset

Results of the AAT building algorithm applied to ICBM 152 dataset is shown of Fig. 1. Comparing the GTC overlap ratios and sharpness for LDD, *s*=1 and *g*=1 were chosen as they produced the best results (Fig. 1a). The same parameters were used to create AAT using the DD algorithm. NDD, LDD and ANTs yield the best GTC overlap. Surprisingly, ANIMAL achieves the highest Sharpness results, followed by ANTs and LDD (Fig 1b). The ATT voxel-by-voxel image intensity standard deviation (SD) can also give an idea of registration quality, where lower SD indicates more consistent registrations. NDD, LDD and DD give the best results for SD (Fig 1c).

**Fig. 1.**
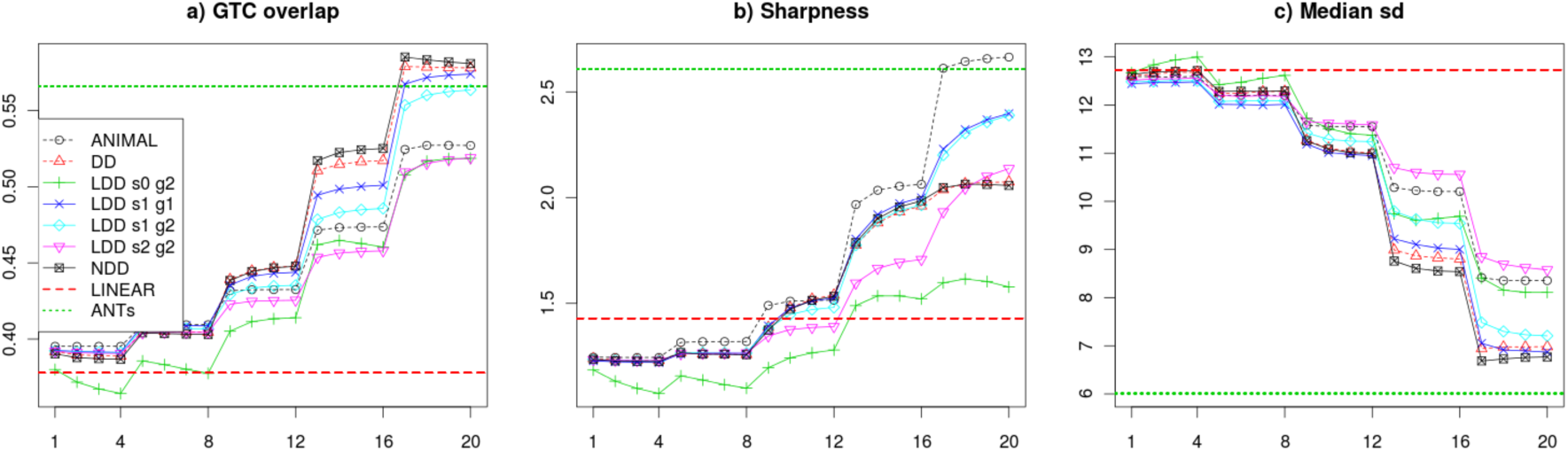
GTC, sharpness and median standard deviation calculated for each iteration of AAT building using the ICBM 152 dataset.

Figure 2 shows one slice of the AATs produced by each algorithm together with voxel-wise GTC overlap ratio map (where higher/brighter voxels indicate better fits between subjects). One can notice that the bulk of the white matter is very well aligned, even when using linear co-registration, however cortex and deep-gray matter structures clearly benefit from using non-linear registration. Also it is worth noting that the cortical folding pattern is slightly different between different methods, whereas deep structures appear very similar.

**Fig. 2.**
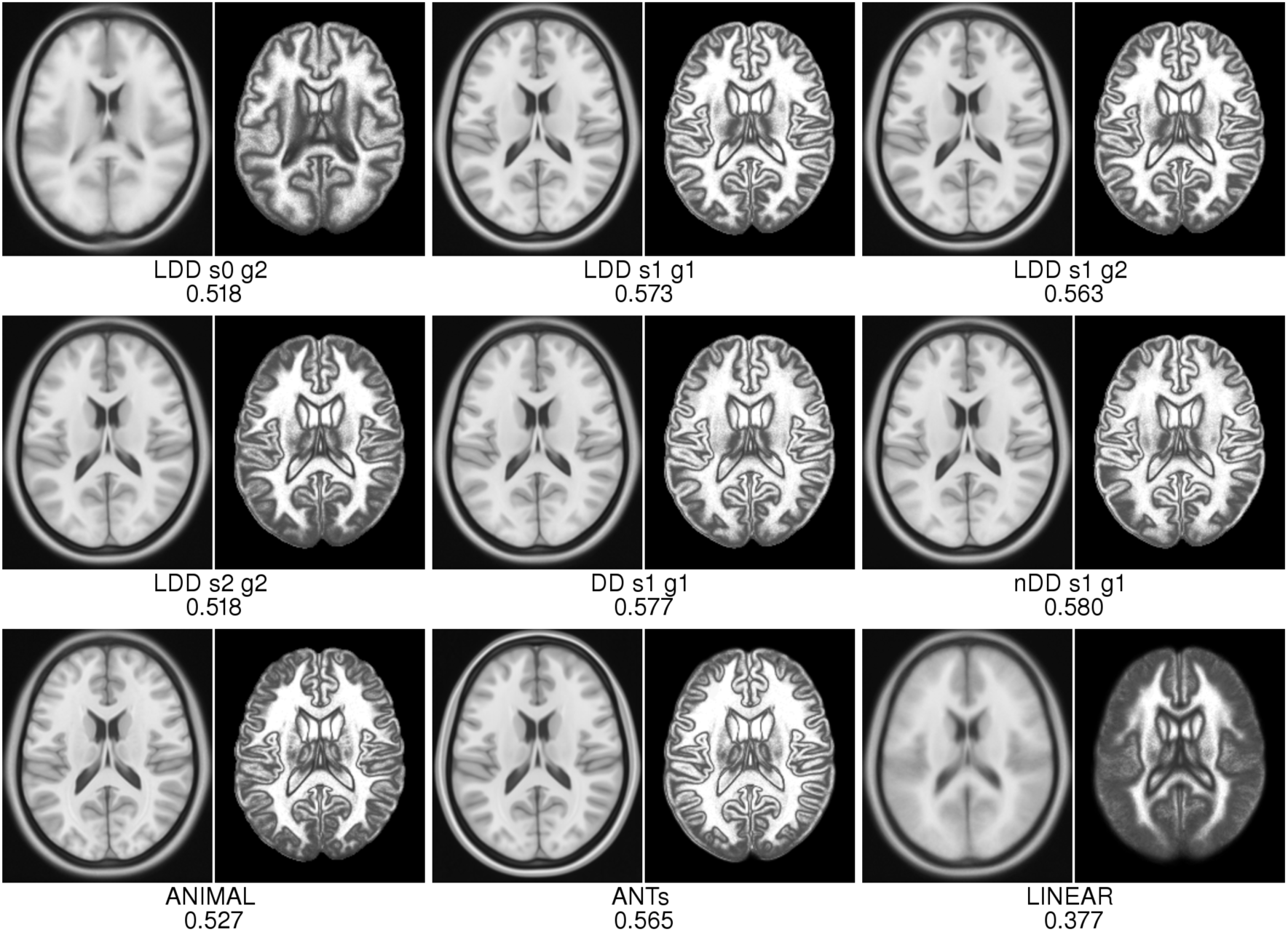
Final results with ICBM 152 dataset. In each pair, the AAT produced by each algorithm (left) is shown with the voxel-wise GTC map (right), where black corresponds to zero overlap and white, 100% overlap. The number corresponds to the overall GTC overlap.

### Results of LPBA40 Average Anatomical Template construction

Figure 3 shows the results of each algorithm on LPBA40 dataset. In contrast to ICBM152 dataset, we have calculated pair-wise GTC overlap ratio between structure label-sets of each subject warped into the space of the AAT, with a total of 780 pairs in each case. The pair-wise tissue GTC results in Fig. 3a show that the three best techniques (in order) are NDD, DD and LDD (p<0.00001, non-parametric Friedman rank sum test). The pair-wise structure GTC results in Fig. 3b show the three best techniques are ANIMAL, NDD and LDD, and while significant, the differences are very small. ANIMAL and ANTs obtain the lowest HE values. Interestingly, while ANTs, ANIMAL and NDD have similar pair-wise structure GTC, the ANTs and ANIMAL HE values are much smaller than NDD, indicating that smoother transforms can achieve similar structure segmentations.

**Fig. 3.**
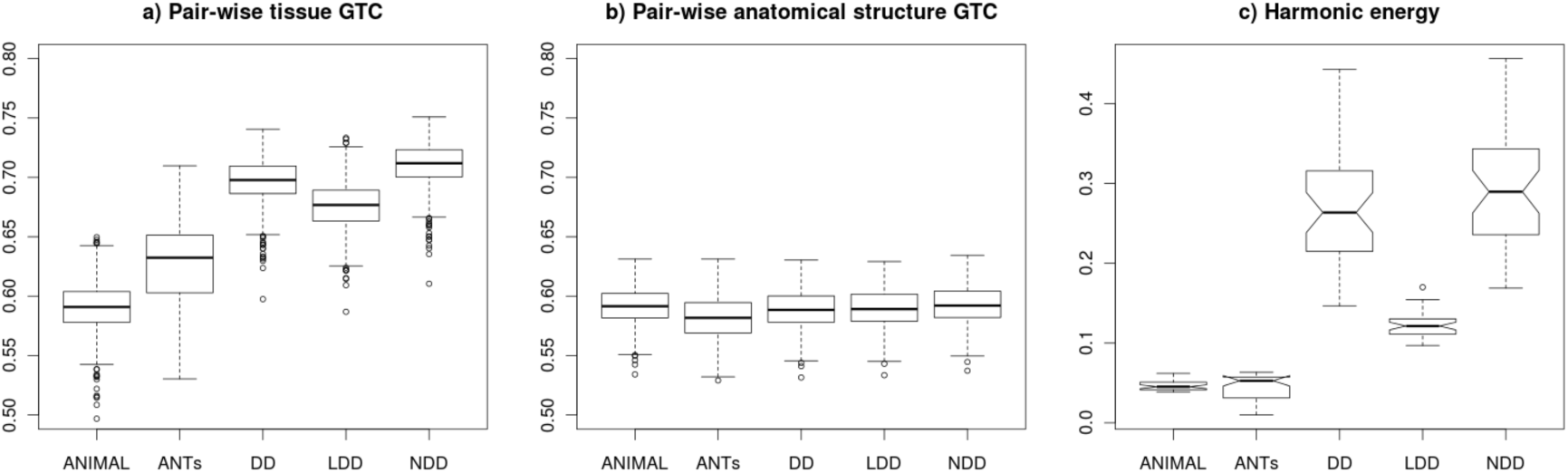
Pair-wise GTC overlap ratio of tissue classes (a), pair-wise overlap ratio of anatomical structures (b) and HE distribution of the deformation fields (c) mapping each subject into the algorithm’s AAT template space.

Figure 4 shows the final AAT and GTC overlap maps produced by each method. Note that LPBA40 dataset contains 55 anatomical structures, but we used only 53; cerebellum and brainstem were excluded since these structures were masked away in the delineation space. Figure 4 demonstrates that the overlap ratio is relatively high at the center of each cortical structure, while the edges do not agree very well for all presented algorithms. This reduction in the overlap metric at structure borders might be caused by both anatomical variability of the human cortex and difficulty of delineating anatomical structures by the human raters.

**Fig. 4.**
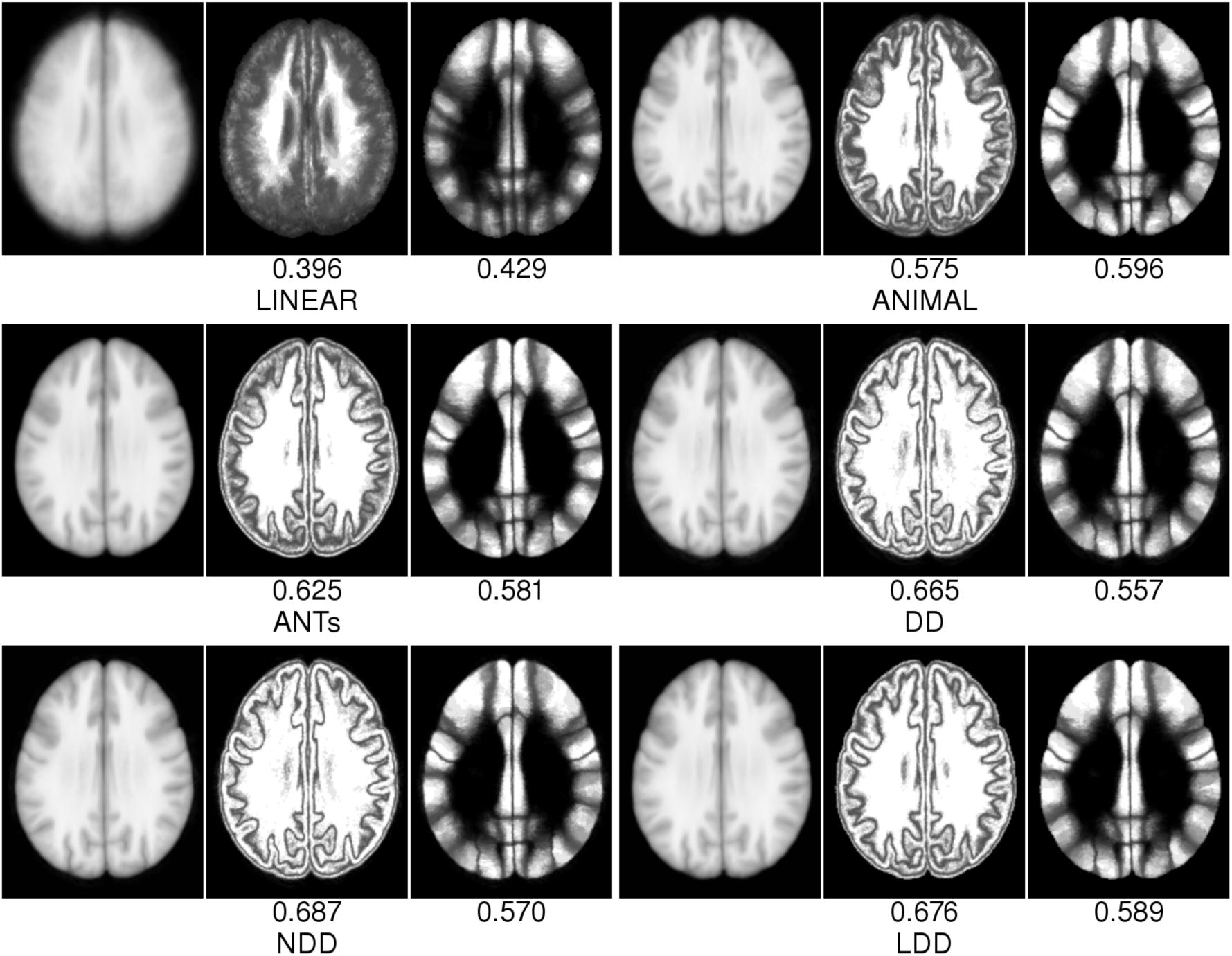
AAT and GTC maps of tissue classes and anatomical structures for each method.

## 4 Discussion and Conclusions

Our experiments demonstrate that the NDD, DD and LDD gave the best tissue-class GTC on both the ICBM152 and LPBA40 databases; ANIMAL achieved the highest Sharpness results, followed by LDD. There is very small difference between algorithms w.r.t. pairwise anatomical structure GTC, while ANIMAL and ANTs yielded the smoothest transforms w.r.t. HE. Our experiments show that application of fully diffeomorphic methods do not automatically guarantee an increase of accuracy in average anatomical template construction, at least w.r.t overlap of tissue classes and anatomical labels, while all techniques yielded well-behaved transformations. In fact, the overall best performing method in our experiments was Thirion’s Demons based algorithm (NDD) with symmetrized gradient calculation (Vercauteren, Pennec et al. 2009).

